# An Invariants-based Method for Efficient Identification of Hybrid Species From Large-scale Genomic Data

**DOI:** 10.1101/034348

**Authors:** Laura S. Kubatko, Julia Chifman

## Abstract

Coalescent-based species tree inference has become widely used in the analysis of genome-scale multilocus and SNP datasets when the goal is inference of a species-level phylogeny. However, numerous evolutionary processes are known to violate the assumptions of a coalescence-only model and complicate inference of the species tree. One such process is hybrid speciation, in which a species shares its ancestry with two distinct species. Although many methods have been proposed to detect hybrid speciation, only a few have considered both hybridization and coalescence in a unified framework, and these are generally limited to the setting in which putative hybrid species must be identified in advance. Here we propose a method that can examine genome-scale data for a large number of taxa and detect those taxa that may have arisen via hybridization, as well as their potential “parental” taxa. The method is based on a model that considers both coalescence and hybridization together, and uses phylogenetic invariants to construct a test that scales well in terms of computational time for both the number of taxa and the amount of sequence data. We test the method using simulated data for up 20 taxa and 100,000bp, and find that the method accurately identifies both recent and ancient hybrid species in less than 30 seconds. We apply the method to two empirical datasets, one composed of *Sistrurus* rattlesnakes for which hybrid speciation is not supported by previous work, and one consisting of several species of *Heliconius* butterflies for which some evidence of hybrid speciation has been previously found.

---

Large-scale genomic data present many challenges in the inference of the evolutionary history of a collection of species. The most notable of these is the development of methods for inferring species-level phylogenetic relationships from multiple gene alignments that simultaneously incorporate the evolutionary processes that are known to contribute to variability in histories for the individual genes. Two important processes are incomplete lineage sorting (ILS) and hybridization (Maddison 1997). ILS results when two gene copies fail to coalesce in the most recent ancestral population and is commonly modeled by the coalescent process, which provides a link between the species tree and the gene trees that represent the phylogenetic history for each gene (Kingman 1982a,b; Tavare 1984). In particular, multispecies coalescent theory models probabilities of rooted gene tree topologies within a given rooted species tree topology and has been used to derive the various probability distributions on gene trees given a particular species tree (Tajima 1983; Takahata and Nei 1985a; Pamilo and Nei 1988; Rosenberg 2002; Rannala and Yang 2003; Degnan and Salter 2005). To date, many methods have been proposed for estimation of species phylogeny from multi-locus data based on the coalescent process (e.g., BEST (Liu and Pearl 2007), *BEAST (Heled and Drummond 2010), STEM (Kubatko et al. 2009), MP-EST (Liu et al. 2010), SNAPP (Bryant et al. 2012), SVDquartets (Chifman and Kubatko 2014) (now implemented in PAUP* (Swofford 1998)), ASTRAL (Mirarb et al. 2014), among others).

Hybridization is another evolutionary process that can cause variability in gene trees within the containing species tree. It generally refers to the interbreeding of individuals from distinct populations, resulting in the production of a hybrid species that shares genetic information with both parental species. Hybridization between distinct species can occur for many generations with fertile offspring, making it possible for a new species to be formed. If the hybridization does not result in the formation of a new lineage, the process is termed introgression or introgressive hybridization (Dowling and DeMarais 1993; Roques et al. 2001; Thorsson et al. 2001; Salzburger et al. 2002; Weigel et al. 2002; Good et al. 2003; Grant et al. 2004; Mallet 2005, 2007; Baack and Rieseberg 2007). Despite the earlier belief that hybridization was rare, numerous recent studies have shown that hybrid speciation occurs in both plants and animals (Rieseberg 1997; Gross and Rieseberg 2005; Buerkle et al. 2000; Bullini 1994; Nolte et al. 2005; DeMarais et al. 1992; Gompert et al. 2006; Schwarz et al. 2005; Mavarez 2006; Meyer et al. 2006; Mallet 2007). Hybridization has been recognized as an important mechanism for the evolution of new species and recent estimates indicate that approximately 25% of plants and 10% of animals hybridize (Seehausen 2004; Mallet 2005, 2007; Baack and Rieseberg 2007; Mallet 2007). However, inference of hybridization cannot be based solely on observed gene tree variability since other processes (e.g., incomplete lineage sorting and gene duplication and loss) may contribute to disagreements in single-gene phylogenies (Maddison 1997).

Several models and methods have been developed to detect hybridization. One group of methods involves the identification and removal of hybrids prior to phylogenetic analysis, with the hybrids added to the inferred tree by connecting them to their parental species (Rieseberg and Morefield 1995; Posada 2002; Gauthier and Lapointe 2007). Joly et al. (2009) developed a method and software (JML; Joly (2012)) for identifying introgressed sequences by proposing that for some hybridization events the minimum distance between two sequences will be smaller than for incomplete lineage sorting. Another test that was originally developed to test ancient admixture is based on a relative abundance of ABBA or BABA single nucleotide patterns that can be evaluated using Patterson’s D-statistic (Green et al. 2010; Durand et al. 2011; Patterson et al. 2012). Meng and Kubatko (2009) proposed a model for detecting hybridization under the coalescent model and used both a maximum likelihood and a Bayesian framework for inference. An extension to that model was later provided by Kubatko (2009) by utilizing gene tree densities for inference. Yu et al. (2014) also proposed a likelihood method that accounts for both reticulate evolutionary events and incomplete lineage sorting by providing methods for computing the likelihood of a phylogenetic network under the coalescent model. This method, as well as some earlier variations of it, is implemented in the software PhyloNet (Than et al. 2008).

In this paper we develop a method for detecting and quantifying the extent of hybridization using a coalescent-based model that is fast and accurate. At the heart of our method are special relations called *phylogenetic invariants*, which are functions (usually polynomials) in the site pattern probabilities that evaluate to zero on any probability distribution that is consistent with the tree topology and associated model. Invariants have been introduced by Cavender and Felsenstein (1987) and Lake (1987) as a means for phylogenetic reconstruction, and have recently been gaining popularity for use in phylogenetic tree inference (Eriksson 2005; Casanellas and Fernández-Sanchez 2011; Chifman and Kubatko 2014). Here we propose using a ratio between two linear invariants in site pattern probabilities to develop statistics that accurately identify hybrid taxa. Because these statistics are functions of site pattern probabilities across multi-locus or SNP data, they can be rapidly computed. In addition, we can derive the mean, variance, and asymptotic distribution of these invariants, enabling development of a hypothesis test for hybridization when the number of sites is large. We begin by giving the theoretical details of our model, and then evaluate the performance of several possible invariants-based statistics for four-taxon trees using simulation. The best-performing of these statistics, which we call the Hils statistic, is then evaluated for larger trees using simulation, with hybridization events at various “depths” of the tree (i.e., hybridization between tip species and hybridization between ancestral species). Finally, we apply our method to several empirical data sets, including the *Sistrurus* rattlesnakes and *Heliconius* butterflies.

## Methods

### A Coalescent-based Model for Hybridization

We consider here the model originally proposed by Meng and Kubatko (2009) in which data arise along a phylogenetic species tree via an evolutionary process that allows for the possibility of both hybridization and incomplete lineage sorting, as modeled by the coalescent process. Hybridization cannot be modeled by a bifurcating phylogenetic tree, thus it is common to represent hybridization on a phylogeny by a horizontal line connecting two lineages of an otherwise-bifurcating phylogeny, as shown in the leftmost panel of Figure 1. This tree represents the evolutionary history of the species as a whole, and depicts a hybrid origin for taxon H. We refer to species *H* as the hybrid species, and to species *P*_1_ and *P*_2_ as the parental species. The times labeled by *τ_i_* are speciation times, and in general we refer to the tree topology *S_γ_* together with its vector of speciation times *τ* by (*S_γ_, τ*). The data arising along this phylogenetic species tree are a collection of site patterns. Letting *X_Y_* ∈ {*A, C,G,T*} denote the nucleotide observed for species *Y* at a specific location in the DNA sequence, we define a site pattern 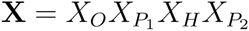 as an assignment of nucleotides to all species. We represent the site pattern probability on the species tree (*S_γ_, τ*) for a particular observation *ijkl* at the tips of the tree by

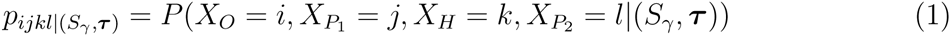

for *i, j, k, l* ∈ {*A, C, G,T*}.

**Figure 1:**
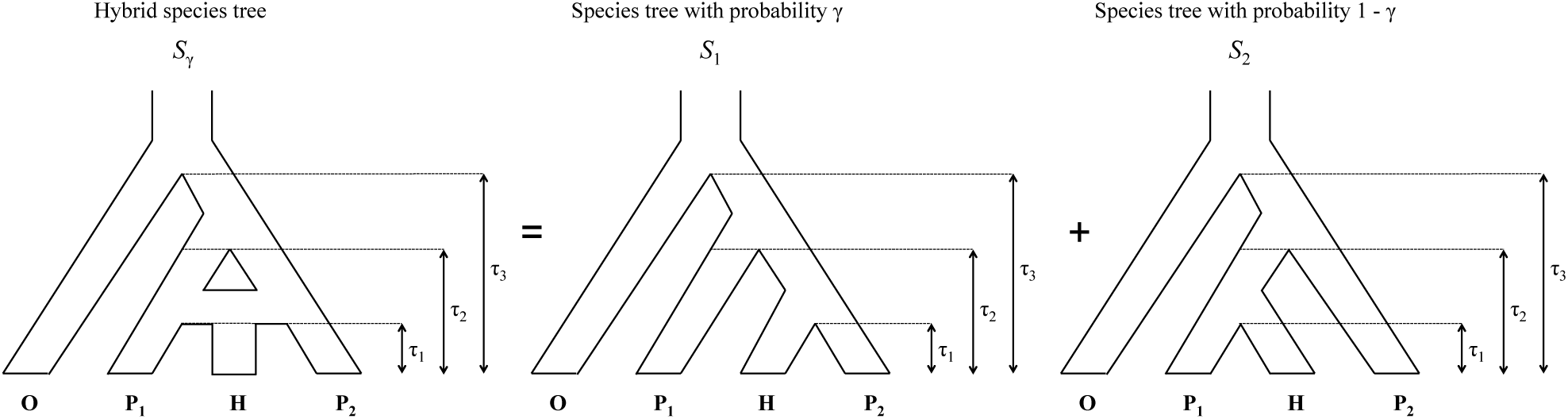
Model for the species-level relationships among four taxa under the coalescent model with hybridization. Here taxon *H* is a hybrid of taxa *P*_1_ and *P*_2_*·*

Our model defines the probability distribution on the space of all 4^4^ = 256 site patterns under a model that allows both ILS and hybridization via a three-stage process. First, the hybrid species is assigned one of its two putative parents, with probability *γ* of selecting parental species *P*_1_ and probability 1 − γ of selecting parental species *P*_2_ (resulting in trees *S*_1_ and *S*_2_ in Figure 1 being the “parental species trees”, respectively). Next, a gene tree is generated along the parental species tree from step 1 through the standard coalescent process (see, e.g., Kingman (1982a,b); Tavaré (1984); Rannala and Yang (2003); Tajima (1983); Takahata and Nei (1985b); Pamilo and Nei (1988); Wakeley (2009)). Finally, a site pattern is generated along the gene tree from step 2 according to one of the standard Markov substitution models (e.g., the GTR+I+Γ model (Lanave et al. 1984) or one of its sub-models). Combining steps 2 and 3, we see that the probability for site pattern *ijkl* for a given species tree *S_i_*, *i* ∈ {1, 2}, is given by

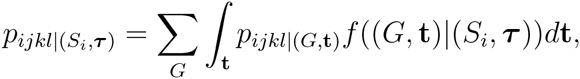

where (*G*, **t**) represents a gene tree with topology *G* and branch lengths **t**, *pijkl|*(*G*, **t**) is the probability of the particular observation *ijkl* at the tips of gene tree (*G*, **t**), and *f*((*G*, **t**)|(*S_i_*, ***τ***)) is the joint density of (*G*, **t**) conditional on the species tree (*S_i_*, ***τ***). A full description of the computations required for this model are given in Chifman and Kubatko (2015), and we do not review them here. Finally, we write the site pattern probability on a hybrid species tree as

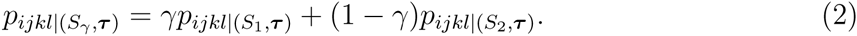

For our purposes, it suffices to view the collection of site patterns observed in an empirical data set as a sample of observations from the probability distribution defined by the {*p_ijkl_*|(*S_γ_*, ***τ***)|*i*, *j, k,l* ∈ {*A,C,G,T*}}. We call data generated in this way “coalescent independent sites” and refer to this model as the “coalescent independent sites model”.

Let *N****_X_*** be the number of sites with site pattern **X** observed in a sample of *N* sites generated from hybrid species tree (*S_γ_*, ***τ***) under this coalescent-with-hybridization model. Define **p** = (*p_AAAA_*|(*S_γ_*, *τ*)*, p_AAAC_*|(*S_γ_*, *τ*),… ,p_TTTT_|(*S_γ_*, *τ*)) and 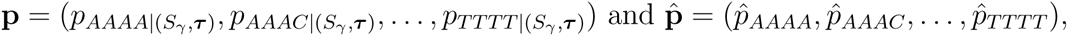, where *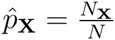*. Then,

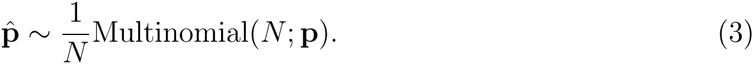

When *N* is large, the 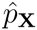 are approximately normally distributed, and thus the sampling distributions of statistics based on the 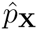 can be derived. We next describe how these ideas can be used to build tests for hybridization.

### Invariants-based Hypothesis Tests for Hybridization

As mentioned in the Introduction, our tests are based on phylogenetic invariants, which are polynomials in the site patterns that evaluate to zero on one tree topology but do not evaluate to zero for at least one tree of a different topology. Consider four linear relationships that arise on the hybrid phylogenetic species tree (*S_γ_*, ***τ***) as described in the previous section:

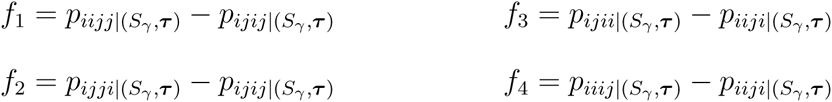

where *i* ≠ *j* ∈ {*A, C, G,T*}. It can be shown that *f*_2_ and *f*_4_ are zero when evaluated on site pattern probabilities that correspond to the species tree *S*_1_, while *f*_1_ and *f*_3_ are non-zero (see Chifman and Kubatko (2015) for details). Similarly, *f*_1_ and *f*_3_ are zero when evaluated on site pattern probabilities that correspond to tree *S*_2_, while *f*_2_ and *f*_4_ are not. However, when the site pattern probabilities correspond to the tree (*S_γ_*, ***τ***) with *γ* ∈ (0,1), none of the four linear relations are zero.

What is special about these functions is that their ratio is a function of *γ* ∈ (0,1):

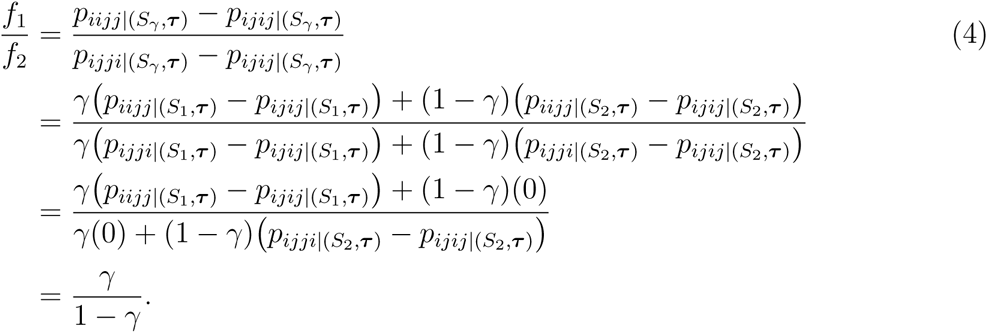

Notice that the last equality holds because *p_ijji_*|(*S*_2_*, τ*) – *p_ijij_|*(*S*_2_*, τ*) = *p_iijj_*|(*S*_1_*, τ*)–*p_ijij_*|(*S*_1_*, τ*).

Using a similar argument we find that

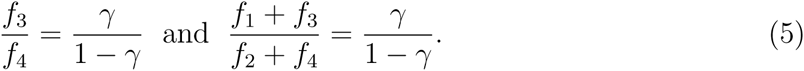

If we consider cumulative site pattern probabilities then the results in Equations (4) and (5) still hold. By a cumulative site pattern we mean, for example, 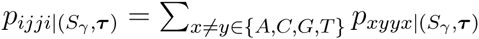. Under the JC69 model (Jukes and Cantor 1969), each of the terms in the sum will have the same value, regardless of the choice of *x* and *y;* under more complex models, these probabilities will vary depending on the particular *x* and *y*. We implement the JC69 version of the test here, though we use simulation to assess the performance under more complicated models.

Using the ratios in Equations (4) and (5) we construct formal significance tests of the following hypotheses:

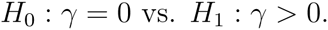

Here we consider the ratio 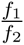 to illustrate the procedure. First, we estimate this ratio using the site pattern probabilities observed in the sample,

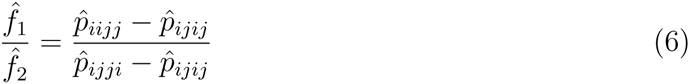

To use this estimator as a test statistic in a hypothesis test, we need the distribution of the statistic when the null hypothesis is true. We first consider distributional results for the numerator and denominator separately. Using standard results for the multinomial distribution, we have

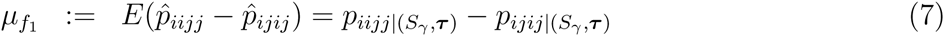

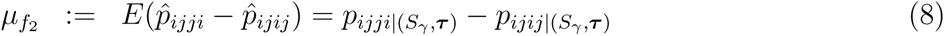

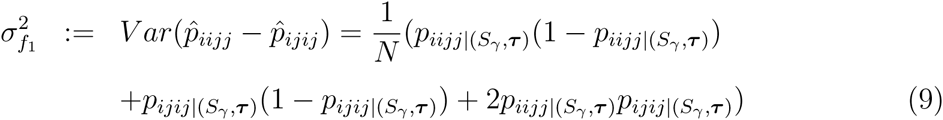

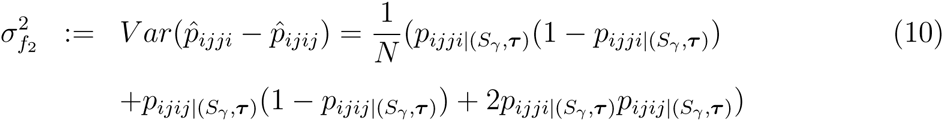

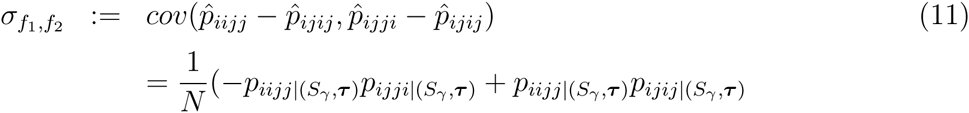

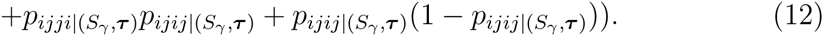

Now, using the fact that when the sample size *N* is large we have 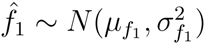 and 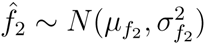, we apply the Geary-Hinkley transformation (Geary 1930; Hinkley 1969) to the ratio 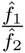 to get

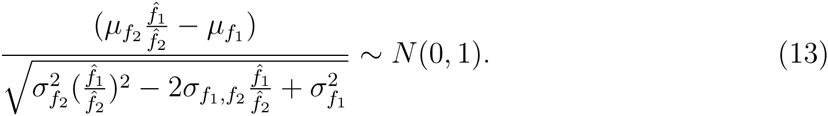

The term on the left-hand side of the above equation depends on several unknown quantities. We estimate these by substituting the observed site pattern frequencies into Equations (7) - (12) and re-arrange the expression in Equation (13) to obtain the test statistic

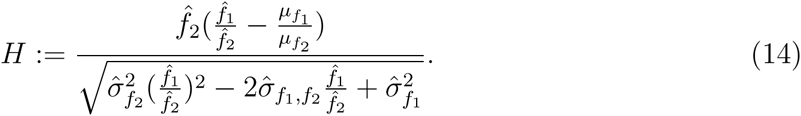

We call the statistic *H* the Hils statistic, in honor of Professor Matthew H. Hils^1^. Under the null hypothesis that *γ* = 0, *H* ~ *N*(0,1) for large *N* with 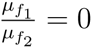, and the hypothesis test can be carried out by comparing the observed value of the test statistic with a standard normal distribution. Tests based on the ratios 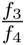 and 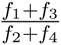 can be derived analogously.

### Extension to Larger Trees

The hypothesis test derived in the previous section deals with the case in which four taxa are specified, with one of the four taxa identified as the putative hybrid species. In many settings, however, primary interest is in searching over a large collection of species with the goal of identifying which species might have arisen via a process that involved hybridization at some point in the past. To address this, we consider a large collection of sequences, and suppose that an outgroup sequence can be identified. For each subset of four sequences consisting of three sequences plus the outgroup, we carry out the above test of hybridization for different assignments of the three ingroup sequences to the hybrid and parental taxa. Of the three possible choices for the hybrid taxon, we consider only two of those, eliminating from consideration the one for which 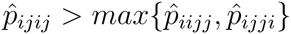, since this implies that the two parental taxa are more closely related than either is to the putative hybrid. For a data set of *n* +1 sequences with one outgroup sequence, this results in 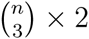 hypothesis tests. To handle the issue of multiple comparisons, we use the Bonferroni correction, which is conservative in this case because the tests are correlated. Thus, if an overall *α*-level test is desired, we report significant evidence of hybridization when the p-value computed for a particular comparison is smaller than 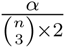.

Our method is implemented in a program written in the C language called HyDe (Hybrid Detection), available at http://www.stat.osu.edu/~lkubatko/software/. The program takes a Phylip-formatted input alignment, with the outgroup sequence specified, and outputs a test statistic for every collection of three species in the input data.

### Simulation Studies

*Four-taxon trees*.— Our first set of simulation studies involves assessing the level and the power of the tests under various choices of the sample size, species trees branch lengths, and value of *γ* for four-taxon trees. We used the program COAL (Degnan and Salter 2005) to simulate gene trees from the two parental species trees in Figure 1 with *γ* values of 0, 0.1, 0.2, 0.3, 0.4, and 0.5 and for two sets of speciation times: *τ*_1_ = 0.25, *τ*_2_ = 0.5, *τ*_3_ = 1.0 (the “short” setting) and *τ*_1_ = 0.5, *τ*_2_ = 1.0, *τ*_3_ = 2.0 (the “long” setting). For each setting, we simulated *N* = 50,000,100,000, 250,000 and 500, 000 coalescent independent sites under the GTR+I+Γ model using Seq-Gen (Rambaut and Grassly 1997)(Seq-Gen options: `-mGTR -r 1.0 0.2 10.0 0.75 3.2 1.6 -f 0.15 0.35 0.15 0.35 -i 0.2 -a 5.0 -g 3)`. For each parameter setting, we generated 500 replicate data sets.

For each simulated data set, we tested the null hypothesis that *γ* = 0 using the test statistics corresponding to the ratios in Equations (4) and (5) at level *α* = 0.05. We estimate the power of each test as the proportion of the 500 replicates for which the null hypothesis was rejected (when γ = 0, this gives an estimate of the level of the test). We also considered using each of the statistics to estimate the true hybridization parameter, γ. We report the mean of the estimated *γ* values, as well as the standard deviation and the mean squared error, for each parameter setting.

*Larger trees*.— To examine the performance of our method for larger taxon samples, we considered trees containing 8 species and an outgroup, and trees containing 19 species and an outgroup. We also considered both recent hybridization and more ancient hybridization in each case. Our model trees are shown in Figure 2. For each model tree, we generated 125 data sets containing 100,000 coalescent independent sites for γ = 0, 0.1, 0.2, 0.3, 0.4, and 0.5 as follows. First, 100000γ gene trees were generated from the species tree formed by connecting the hybrid taxon to the “left” parental lineage, and 100000(1 – γ) gene trees were generated from the species tree formed by connecting the hybrid taxon to the “right” parental lineage. For each gene tree, one coalescent independent site was generated using Seq-Gen (Rambaut and Grassly 1997) under the GTR+I+Γ model (Seq-Gen options: `-mGTR -r 1.0 0.2 10.0 0.75 3.2 1.6 -f 0.15 0.35 0.15 0.35 -i 0.2 -a 5.0 -g 3)`. Each simulated data set was then given to our program with the outgroup specified, and the Hils statistic was computed for each possible combination of parents and hybrids. A cut-off for significance was determined using a Bonferroni correction with base level *α* = 0.05, and the putative hybrid and parents were reported for any statistic whose p-value fell below *α/M*, where M was the total number of comparisons. We summarized results by counting the number of “True Positives” (data sets for which the correct hybrid and parental taxa are correctly identified), “Correct Sets” (data sets for which the correct hybrid and parental taxa are identified, but their assignment to which is the hybrid and which are the parental taxa is ambiguous), and “False Positives” (data sets for which an incorrect set of taxa are identified as being subject to hybridization).

**Figure 2:**
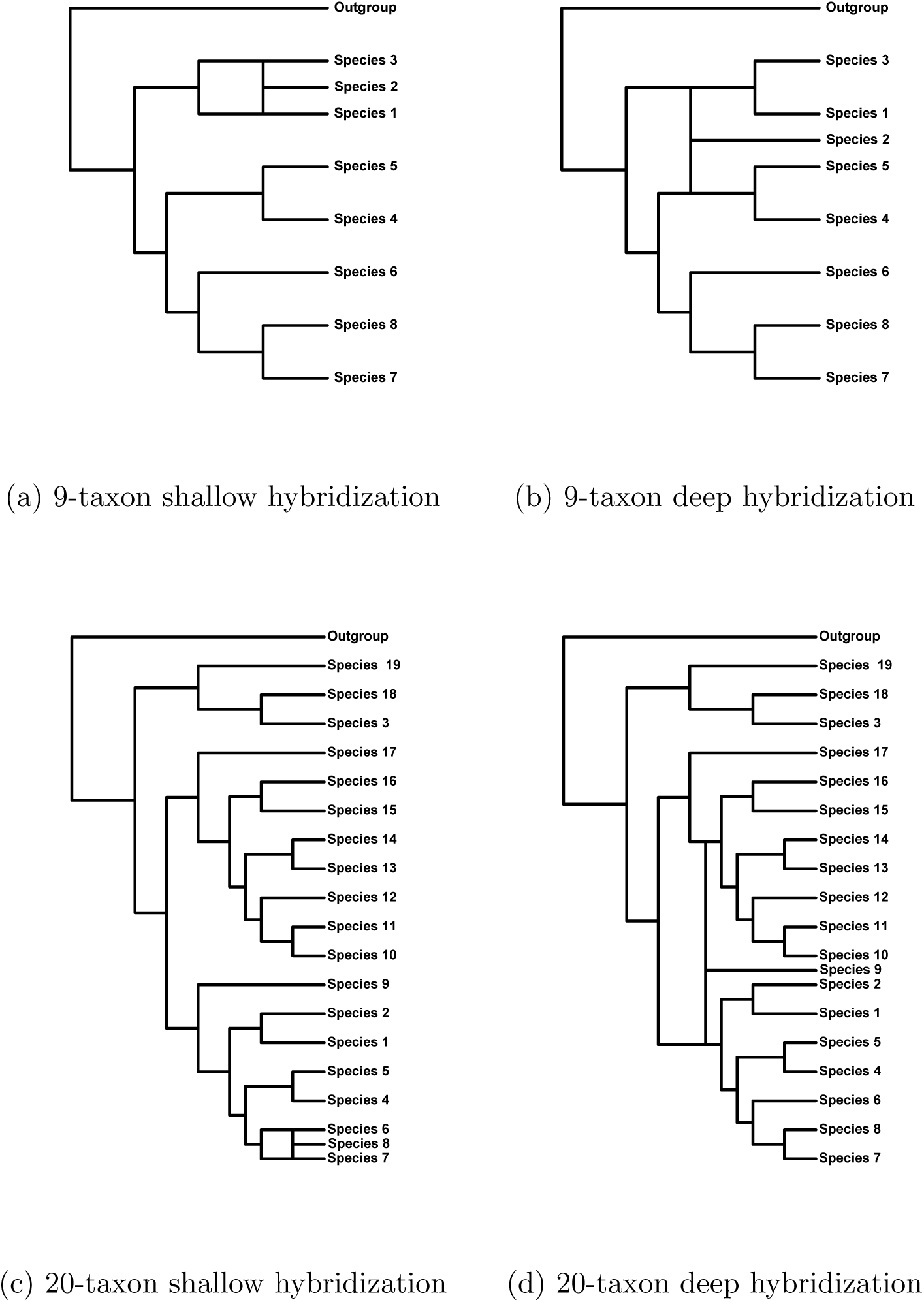
Model trees with 9 and 20 taxa and with either shallow or deep hybridization used for the simulation studies.

Because many of the genome-scale datasets being generated today are multilocus datasets (rather than being generated under the coalescent independent sites model used here), we also simulated data under multilocus models. These simulations proceeded exactly as described above, except that rather than simulating 100,000 coalescent independent sites, we simulated 1,000 genes each of length 100bp. This choice was made to mimick the short read lengths generated by next-gen sequencing methods. We summarized these results in the same manner as described above. We justify application of our methodology to multilocus data in the Discussion section.

*Empirical examples*.— We have also explored the performance of our method on two empirical data sets; the *Sistrurus* rattlesnakes and *Heliconius* butterflies. The *Sistrurus* rattlesnakes are found across North America and are currently classified into two species, *Sistrurus catenatus* and *S. miliarius*, each with three putative subspecies. The dataset consists of 19 genes sampled from 26 rattlesnakes: 18 individuals within the species *Sistrurus catenatus* (with subspecies *S. c. catenatus* (Sca, 9 individuals), *S. c. edwardsii* (Sce, 4 individuals), and *S. c. tergeminus* (Sct, 5 individuals)); six within species *Sistrurus miliarius* (with subspecies *S. m. miliarius* (Smm, 1 individual), *S. m. barbouri* (Smb, 3 individuals), and *S. m. streckeri* (Sms, 2 individuals)); and two outgroup species, *Agkistrodon contortrix* and *A. piscivorus*. These data were originally analyzed by Kubatko et al. (2011) to determine species-level phylogenetic relationships. Prior to this analysis, the sequences were computationally phased, resulting in 52 sequences and 8,466 aligned nucleotide positions (data are available at TreeBase ID 11174). These data have been subsequently reanalyzed in several ways. For example, Chifman and Kubatko (2014) used different methodology to infer the species phylogeny, and found agreement with the original analysis of Kubatko et al. (2011). Gerard et al. (2011) used a subset of the data to examine whether several specimens collected in Missouri and assigned to subspecies *S. c. catenatus* were actually hybrid species. They did not find evidence of hybridization, in agreement with other results using different data (Gibbs et al. 2011).

The *Heliconius* butterflies are a diverse group tropical butterflies in the family *Heliconii* that are found throughout the southern United States and in Central and South America. We consider the study of Martin et al. (2013) in which genome-scale data for 31 individuals from seven distinct species were collected and evidence for gene flow between various species was assessed. We examine a subset of these data consisting of four individuals from each of the species *Heliconius cydno, H. melpomene rosina*, and *H. m. melpomene*, as well as one individual from the outgroup species *H. hecale*. Martin et al. (2013) found evidence that *H. m. rosina* is a hybrid of *H. m. melpomene* and *H. cydno*. We obtained the aligned genome-wide data from the complete study of Martin et al. (2013) from Dryad (http://datadryad.Org/resource/doi:10.5061/dryad.dk712) (Martin et al. 2013a), and extracted the 13 sequences of interest. The resulting aligned sequences consisted of 248,822,400 base pairs.

## Results

*Four-taxon simulation studies*.— The results of the four-taxon simulation studies are shown in Figure 3 and Table 1. Figure 3 shows that the various tests behave as we might expect in several ways. First, in all of the cases considered, the power increases as the sample size increases, reaching near 100% when alignments of length 500,000bp were used for many of the simulation conditions. Second, we note that as the value of *γ* increases from 0 (no hybridization) to 0.5 (equal contribution from both parental species), the power to detect hybridization increases as well, with near 100% power for the “long” branch length setting when *γ* ≥ 0.3 for all three of the tests considered. Third, we note that all of the tests are more powerful for data simulated under the “long” branch length setting (Figure 3 (d), (e), and (f)) than for data generated under the “short” branch length setting (Figure 3 (a), (b), and (c)). Finally, we note that all tests appear to achieve the nominal 0.05 level when data are simulated under the null hypothesis (γ = 0).

**Figure 3:**
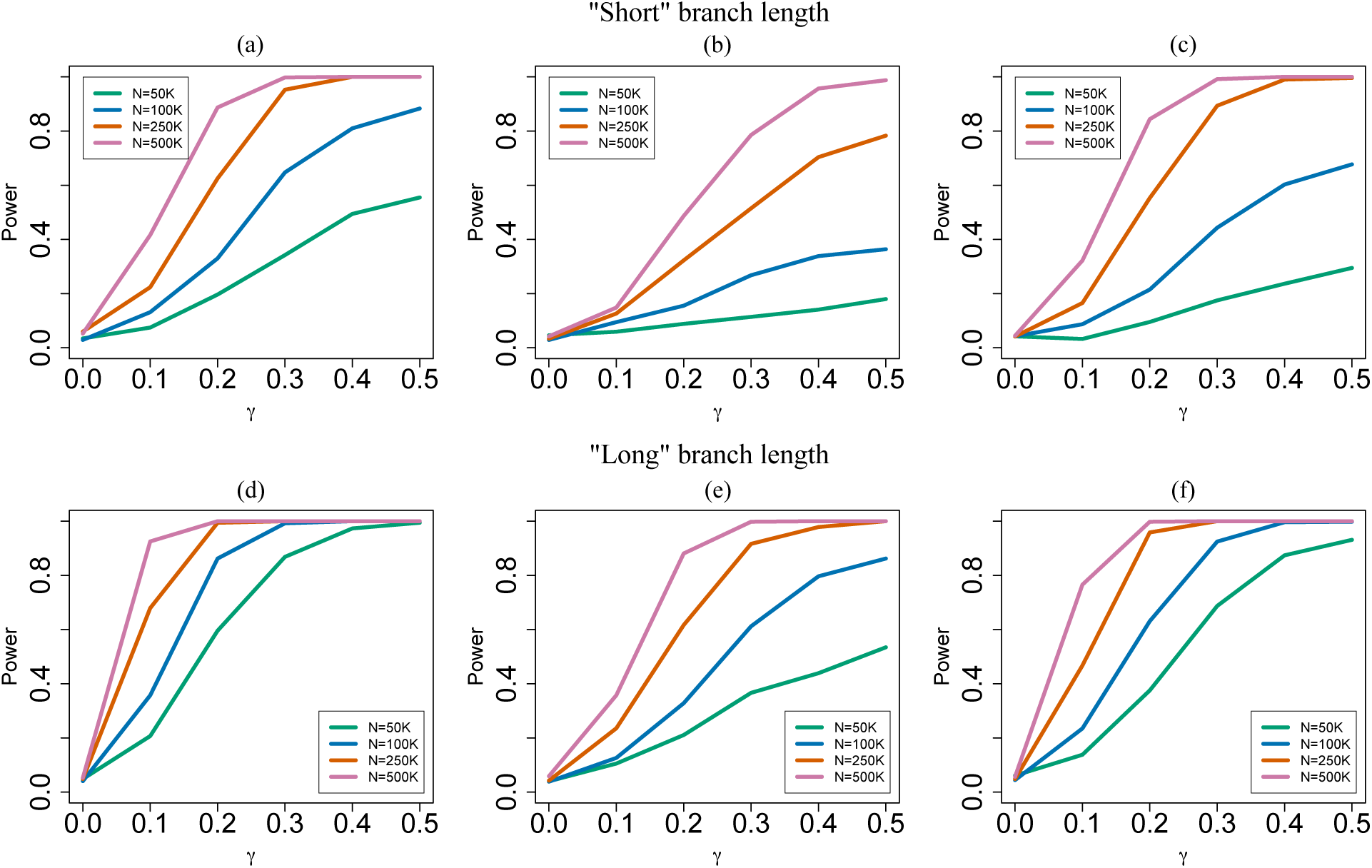
Results of the power simulations for the four-taxon hybrid species tree in Figure 1. Plots (a), (b), and (c) correspond to data simulated for the “short” branch length setting, and ; plots (d), (e), and (f) correspond to data simulated for the “long” branch length settings. Plots (a) and (d) give results for the test based on 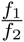; plots (b) and (e) give results for the test based 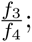; and plots (c) and (f) give results for the test based on 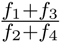.

**Table 1:**
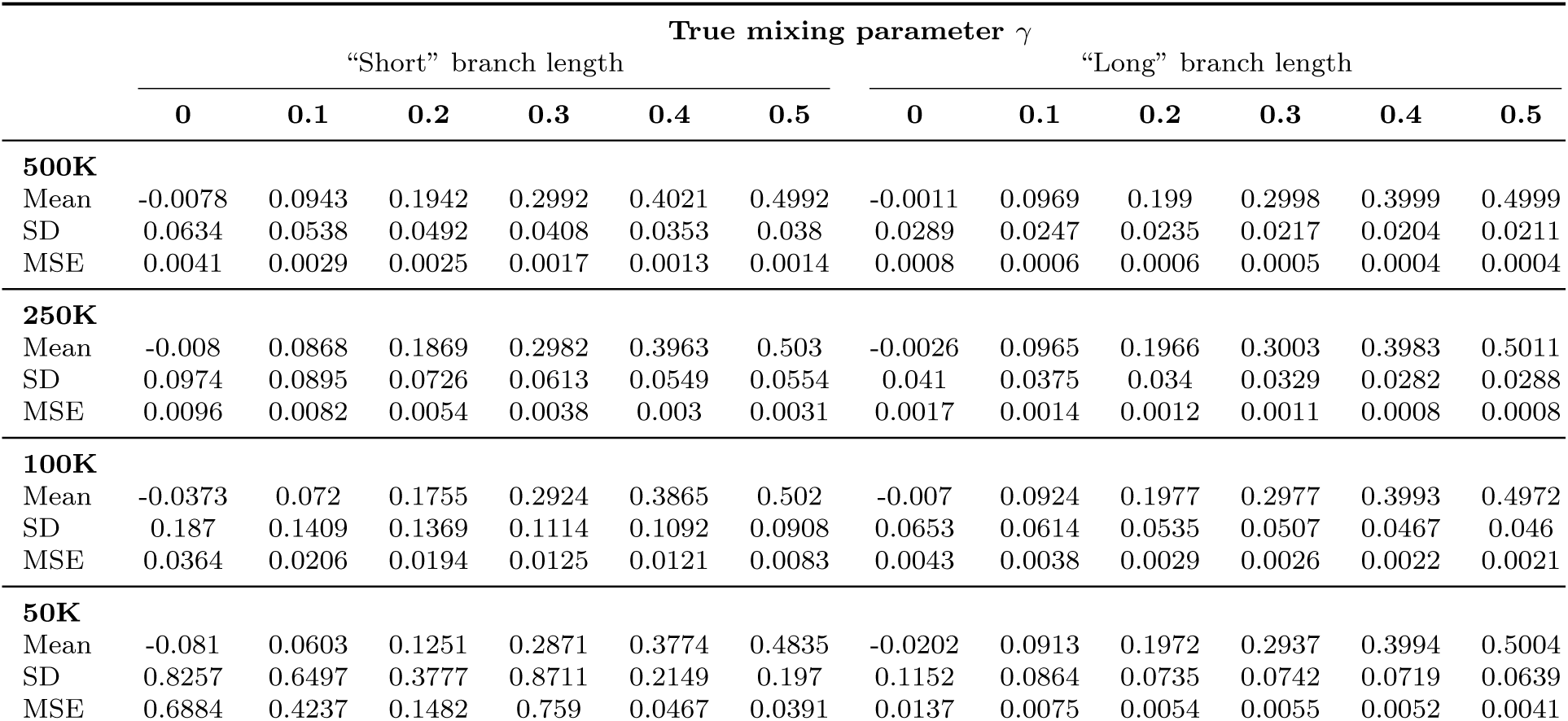
Estimates of the parameter γ using the ratio 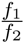 for data simulated on the four-taxon hybrid species tree in Figure 1 with the “short” and “long” branch length settings.

One unexpected result of the simulations designed to address the power was that the test based on 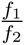 is more powerful than the tests based 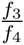 and 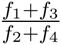. This is most likely due to the variance associated with estimating the various site pattern probabilities that contribute to each invariant. We return to this point in the discussion. Based on this observation, we report results for only the ratio 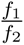 in what follows.

Table 1 gives the results of the four-taxon simulation studies designed to estimate using the ratio 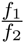. These results also match our intuition about how the method should perform. As the sample size increases, the estimates become closer to the true values used to generate the data, and the variance decreases as the sample size increases. In general, the estimates obtained from the “long” branch length setting are slightly better than those obtained from data generated under the “short” branch length setting. Overall, the method seems to provide very reasonable estimates of *γ*.

*Simulation studies for larger trees*.— The results of the simulation studies for larger trees are given in Tables 2 and 3. For the 9-taxon simulations, we note first that for data generated under the coalescent independent sites model, when *γ* = 0 approximately 5% of the data sets give significant results, and thus the test appears to attain the desired significance level in this case. For the multilocus data sets, however, the type I error rate is larger than the specified 0.05 level, and thus the test appears to reject the null hypothesis more often than it should. When *γ >* 0, we see that the test is powerful for both the shallow and the deep hybridization events and for both types of data, with the power above 90% in both cases when *γ* ≥ 0.2. Furthermore, the test almost always selects the correct assignment of hybrid and parental taxa, with the proportion of times that this is exclusively generated increasing toward 100% as *γ* increases for the coalescent independent sites data. One observation we made that is not reflected in the results in Table 2 is that for data simulated from the tree involving the deep hybridization event, many sets appear as significant when some true relationship is detected. For example, it is common to have the hybrid correctly assigned, but the parental species assigned as belonging to a taxon from the sister clade of the true parent. This is especially true for the multilocus data sets with the deep hybridization event. In other words, this test is good at picking out the hybrid taxon, but not as good at unambiguously picking out its parents when the hybridization event occurs deeper in the tree. This was not the case for the shallow event, where it often got exactly the correct relationships and only those in most cases.

**Table 2:**
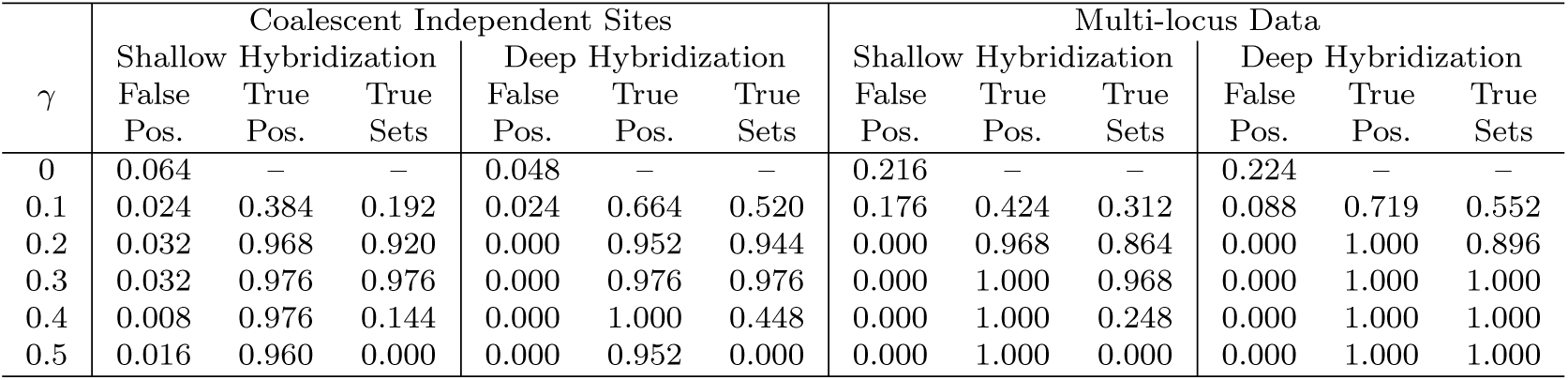
Results of the simulation study for 9 taxa. The columns labeled “False Pos.” refer to the proportion of data sets for which a triplet of taxa were incorrectly identified as involving a hybridization event (false positives); the columns labeled “True Pos.” refer to the proportion of data sets for which the correct triplet of taxa involving the hybridization event was identified *and* the hybrid taxon was correctly identified (true positives); and the columns labeled “True Sets” refer to the proportion of data sets for which the correct triplet of taxa was identified but the hybrid taxa was specified incorrectly. All data sets for which the true set was identified also identified the triplet with the correct hybrid assignment, and thus this proportion is always a fraction of the proportion of true positives.

**Table 3:**
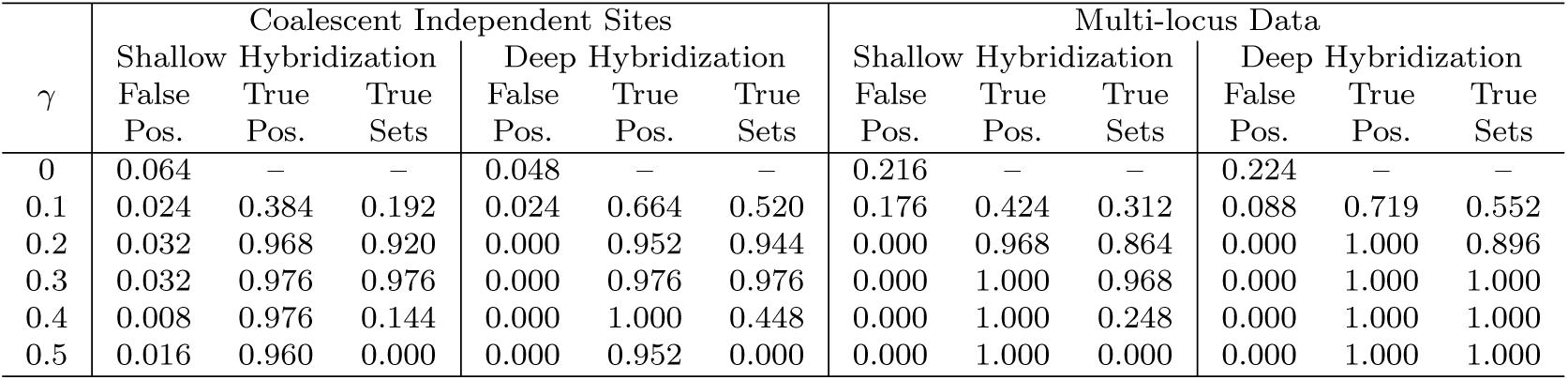
Results of the simulation study for 20 taxa. Column headings are as in Table 2.

The results for the 20-taxon trees are largely the same. The test still demonstrates good power to detect the hybridization event, though the power does not rise above 90% for all settings until *γ* ≥ 0.3, rather than 0.2 as in the 9-taxon case. In addition, the proportion of data sets with “Correct Sets” decreases for the shallow hybridization events in this case, meaning that when a hybridization event is identified, it nearly always involved correct identification of which species was the hybrid and which were the parental species. Though there is a hint of an elevated type I error rate when multilocus data were simulated, the problem is not as dramatic as in the 9-taxon case. Overall, the method maintains its good ability to detect hybrid species.

*Empirical data: Sistrurus rattlesnakes*.— Recall that this dataset contains two species, each containing three subspecies, as well as two outgroup species, for a total of eight tips in the species phylogeny of interest. When analyzing empirical data of this nature, for which several individuals are sampled within each species, our main interest will be in detecting individuals that show evidence of hybrid origin from parental individuals that are members of two different species. The current version of our software will output the test statistic for all assignments of hybrid and parental taxa for a given outgroup, but this output can easily be examined to consider only the comparisons of interest. For the rattlesnake data for a particular choice of outgroup, we can consider all choices of one individual allele from each of three subspecies, and for each such choice, one individual will be assigned to be the hybrid and the other two assigned to be the parental taxa. For example, we can select one Sca individual, one Sce individual, and one Sct individual, and carry out the Hils test for each possible choice of hybrid among these three. Thus, for our particular data set consisting of 18 Sca alleles, 8 Sce alleles, 10 Sct alleles, 2 Smm alleles, 6 Smb alleles, and 4 Sms alleles, there will be 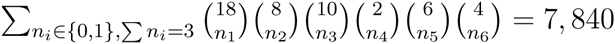 possible choices of three alleles, and two test statistics will be computed on each, resulting in 2 * 7840 = 15, 680 possible comparisons for each choice of outgroup sequence. We carry out the Bonferroni correction within the analysis for each outgroup, and thus each comparison uses significance level *α* = 0.05/15680 = 0.0000032.

An additional practical issue that arose with our empirical data but was not observed with simulated data was that for some choices of three alleles, one or more of the site pattern frequencies *p_iijj_, p_ijij_*, and *p_ijji_* was observed to be 0. To correct for this, we added a small count (0.005) to each observed site pattern count in all cases before computing estimated site pattern frequencies and carrying out the test. With this modification, we find no evidence of hybrid origin for any of the sequences with any choice of outgroup sequence, consistent with other analyses in this group (Gerard et al. 2011; Gibbs et al. 2011).

*Empirical data: Heliconius butterflies*.— This dataset consists of 3 species with 4 individuals sampled per species, plus an outgroup. Thus, the number of comparisons of interest is 4 * 4 * 4 * 2 = 128 and the Bonferroni-corrected level of the tests is 0.05/128 = 0.00039. The analysis of all possible hybrid/parental combinations for the alignment of length ≈ 248 million bp took 16 minutes on a 2x Quad Core Xeon E5520 / 2.26GHz / 32GB desktop linux machine. All comparisons were statistically significant at the 0.00039 level. This result is not surprising, given the previous evidence of hybridization as described in Martin et al. (2013), and given the large sample size. What is interesting, however, is the strength of the evidence for hybridization. For example, across all comparisons in which an *H. m. rosina* individual was specified as the hybrid, the smallest test statistic was 172.6143, indicating overwhelming evidence for hybridization (recall that we are comparing to a standard normal distribution). In contrast, when one of the other species was identified as the hybrid and *H. m. rosina* was (incorrectly) identified as a parental taxon, the values of the test statistic ranged from ~ 55 to 76, again indicating strongly significant deviation from the expected patterns under no gene flow, but not as strong as the case in which the hybrid is correctly identified as *H. m. rosina*. Overall, these results are in agreement with the work of Martin et al. (2013) on this group, and demonstrate the utility of our method in rapidly identifying hybrid taxa from genome-scale data.

## Discussion

We have proposed a method for detecting hybrid species using a model of hybrid speciation that incorporates coalescent stochasticity. The test is based on observed site pattern frequencies, which leads to several convenient properties. First, the computations required for the test can be carried out very rapidly, as all that is required is to obtain counts of observed site pattern frequencies for four taxa of interest. This computation is so rapid that there are essentially no limits on the length of sequences that can be handled by the method, and it is thus appropriate for genome-scale data. Second, observed site pattern frequencies arise from a multinomial distribution under the coalescent hybridization model used here, which allows derivation of the asymptotic distribution of the estimators of the site pattern frequencies. This ultimately leads to a null distribution for testing the hypothesis of interest that is asymptotically normally distributed which provides a straightforward test of the hypothesis of interest. Finally, we note that our method is derived under the assumption that each site has its own underlying gene tree, an experimental design that we propose calling “coalescent independent sites”. The method is thus clearly appropriate for genome-wide SNP data, whether biallelic or not. We argue that the method is also appropriate for multilocus data, in that as the number of loci becomes large and provided that alignment lengths are not biased toward certain gene tree topologies, the proportion of sites observed from a particular gene tree will approach the proportion expected under the coalescent independent sites model. We thus carry out simulations for*f* both multilocus and coalescent independent sites data, and we test our method on an empirical multilocus dataset.

Our simulations show that the method is powerful for detecting hybridization for both recent and ancient hybridization events, although for ancient hybridization events it may be more difficult to pinpoint the precise parental species for the detected hybrids. In addition, the proportional contribution of the two parental species to the genome of the hybrid species can be estimated accurately and unbiasedly. The simulations also show that the method scales extremely well: for 20-taxon trees with 100,000 sites, computations can be completed in less than 30 seconds, while for a dataset with 13 sequences and over 248 million sites, the analysis took less than 20 minutes on an older desktop linux machine. To the extent of our knowledge, this method is thus the only technique available for exploratory hybrid identification for large numbers of sequences using genome-scale data.

The method is based on phylogenetic invariants, and we note that the particular choice of invariants used here was somewhat arbitrary. Indeed, the ABBA-BABA test (Green et al. 2010; Durand et al. 2011; Patterson et al. 2012) is based on the difference of ABBA and BABA patterns similar to our invariant *f*_2_ and it too has been shown to be useful in detecting hybridization. However their statistic is normalized by the total number of observations whereas our method is based on the ratio of two linear invariants leading to a function that d*γ*is γ. Based on this crucial observation we were able to derive Hils statistic for accurate detection of hybridization. We have also noticed that the ratio between*f*_3_ and *f*_4_ was not as powerful, thus it is possible that other invariants may be identified that work as well or better than the ones we have chosen here. It is also possible that invariants that operate on more than four taxa at a time could be determined, with potential improvements in the localization of hybrid and parental taxa for more ancient hybridization events. It is also possible that a set of linear invariants specific to species trees under the coalescent exists and can be classified, and if such a set exists, these species invariants may improve the performance. We suggest that exploring these directions is appealing, as site pattern-based methods provide the possibility of both rapid computation and convenient asymptotic distributions, making them suitable for processing the large genome-scale datasets that are becoming increasingly available. In fact, the performance of these methods improves with sequence length, since site pattern probabilities can be more accurately estimated, with little associated computational cost.

## Footnotes

1 Matthew H. Hils was a Professor of Biology at Hiram College until his untimely death due to cancer in June 2014. He served as academic advisor and research mentor to L.K. during her undergraduate studies, and contributed to her decision to pursue interdisciplinary graduate study tied to the biological sciences. See http://news.hiram.edu/?p=10502.

## Acknowledgements

This work was supported in part by the National Science Foundation under award DMS-1106706 (J.C., L.K.) and NIH Cancer Biology Training Grant T32-CA079448 at Wake Forest School of Medicine (J.C.).

